# Inbreeding depression causes reduced fecundity in Golden Retrievers

**DOI:** 10.1101/554592

**Authors:** Erin T. Chu, Missy J. Simpson, Kelly Diehl, Rodney L. Page, Aaron J. Sams, Adam R. Boyko

**Author notes:** These authors contributed equally to this manuscript (Aaron J. Sams, Adam R. Boyko).

## Abstract

Inbreeding depression has been demonstrated to impact vital rates, productivity, and performance in many domestic species. Many in the field have demonstrated the value of genomic measures of inbreeding compared to pedigree-based estimates of inbreeding; further, standardized, high-quality phenotype data on all individuals is invaluable for longitudinal analyses of a study cohort. We compared measures of reproductive fitness in a small cohort of Golden Retrievers enrolled in the Golden Retriever Lifetime Study (GRLS) to a genomic measurement of inbreeding, F_ROH_. We demonstrate a statistically significant negative correlation between fecundity and F_ROH_.This work sets the stage for larger scale analyses to investigate genomic regions associated with fecundity and other measures of fitness.

## INTRODUCTION

The term “inbreeding depression” encompasses a reduction of a trait, often associated with lifetime fitness, as a sequelae to a sustained rate of breeding of closely related individuals (reviewed in Charlesworth and Willis, 2009; Hedrick and Garcia-Dorado, 2016). While inbreeding depression has been extensively explored in plants (Lande and Schemske, 1984), geographically isolated wild animal populations (Furlan et al, 2012, Hagenblad et al, 2009), and endangered and zoo populations (Roelke et al, 1993), much research of late has addressed the same phenomenon in domestic species, many of which have been selectively bred for performance, production, and companionship. The correlation between inbreeding and impaired production in the dairy, wool, and meat industry has been well described (Ercanbrak et al, 1991; Noren et al, 2015; Mokhtari et al, 2014; Pereira et al, 2017; Perez et al, 2018). More recently, inbreeding has been correlated with reduced performance in Australian Thoroughbred horses (Todd et al, 2018).

In the past, the estimation of inbreeding has relied on in-depth pedigrees, whereby a coefficient of inbreeding (COI), estimated from pedigree-based relationships between ancestors (F_PED_), is used in lieu of measurement of true autozygosity (Wright, 1922). Genomic measures of the COI based on runs of homozygosity (F_ROH_) preclude the need for pedigree-based COIs, which depend heavily on pedigree depth and accuracy (Zhang et al, 2015); even with detailed pedigrees, estimated COIs can deviate substantially from true autozygosity due to recombination and segregation (Hill et al, 2011; Keller et al, 2011). With the availability of high-density SNP arrays and affordable DNA sequencing, F_ROH_ has proven more effective than pedigrees (Huisman et al, 2016) or limited microsatellite panels (Hoffman et al, 2014) in assessing inbreeding and fitness in animal and human populations (Bruniche-Olsen et al, 2018).

As accurate as genome-wide assessments of inbreeding have proven, equally high-quality phenotype data are necessary to detect inbreeding depression. In humans and wild populations, inbreeding depression can be assessed by tracking vital rates--birth rate, mortality rate--in a population over time (Bittles et al, 2002; Robert et al, 2005; Robert et al, 2009; Johnson et al, 2011). In domestic species, additional measures of inbreeding depression include litter size, reproductive success, body size, and performance traits are used (as discussed earlier). Naturally, these analyses can be clouded by external factors including environment, demographics, record completeness and accessibility, and genetic heterogeneity. In that specific regard, the domestic dog, *Canis familiaris*, is an ideal candidate species in which to assess inbreeding depression. In effect, purebred dogs represent naturally occurring populations with limited genetic variation, the result of closed breed registries and strict breed standards for appearance and affect. Further, dogs have an average gestational period of two months and are polytocous, providing rapid collection of fecundity data, and have an average lifespan of roughly 10% of the average human lifespan, permitting timely collection of multigenerational mortality data.

Initiatives for banking of biological samples in combination with standardized, detailed phenotype data are gaining greater traction in the canine community as a means to identify genetic, epigenetic, and environmental variants that impact canine health and longevity. One such initiative, the Morris Animal Foundation’s (MAF) Golden Retriever Lifetime Study (GRLS), seeks to identify genetic and environmental variables that impact longevity in the Golden Retriever (Guy et al, 2015). Known for its sunny coat and disposition, the Golden Retriever is widely recognized as one of America’s favorite dog breeds and is consistently ranked in the top-five highest breeds in AKC registrations annually (American Kennel Club, a). Unfortunately, Golden Retrievers are also overrepresented in neoplasia cases, with more documented mortalities due to cancer than nearly any other breed (Kent et al, 2018; Dobson et al, 2014). And while some genetic variants have been associated with increased risk for certain cancers (Arendt et al, 2015), other major genetic contributors to Golden Retriever lifespan and fitness remain unidentified.

In four years, the GRLS has amassed a sample set of over 3,000 Golden Retrievers, complete with annual biological samples and standardized phenotype data collection from owners and veterinarians (Simpson et al, 2017), and represents a one-of-a-kind dataset for genomic analysis. Here, we combine detailed reproductive data gathered on 93 GRLS participants with high-density SNP genotyping. We evaluate the correlation of the genomic coefficient of inbreeding, F_ROH_, with various indicators of female reproductive success, and we identify a negative correlation between F_ROH_ and live litter size.

## RESULTS

Study participants were drawn from the GRLS cohort of 3,044 dogs. 1,504 were female; 239 of these had been bred at least once. A random stratified sample of 100 dogs, termed the Embark-GRLS cohort, was selected based on number of attempted breedings to enrich for dogs who had been bred several times and had the potential of producing several litters (Summarizing statistics available in Table S1). 93 dogs were successfully genotyped, ranging from 1 to 7 years of age. A total of 407 heats were recorded; heat frequency ranged from 0 to 4 heats per dog per year. Recorded heats for dogs over the age of five years decreased dramatically, likely reflecting the relative youth of the GRLS cohort as well as increased likelihood for elective spay in older bitches. 66 dogs had produced at least one litter, with a total of 99 litters observed. F_ROH_ ranged from 0.187 to 0.479, with mean F_ROH_ of 0.316 (Fig 1).

**Fig 1.**
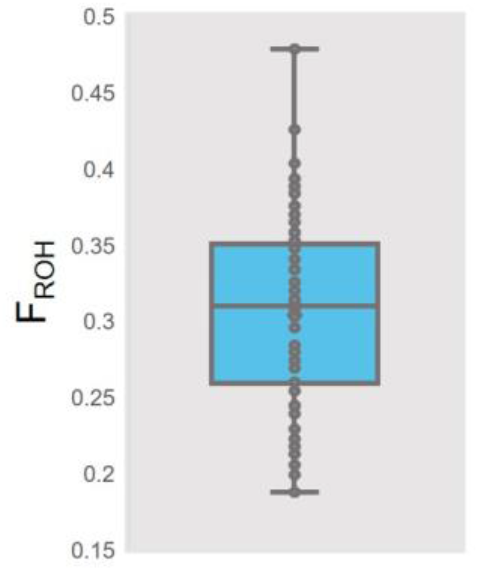
Box and whisker plot of F_ROH_ for 93 genotyped dogs in the Embark-GRLS cohort. F_ROH_ ranged from 0.187 to 0.479, with mean F_ROH_ of 0.316.

Many have demonstrated a negative impact of F_ROH_ on body size (Fredrickson et al, 2012; Lacy et al, 2013; Fareed et al, 2014; Cecci et al, 2018). We regressed the median shoulder measurement for each dog against F_ROH_ and found that in this dataset, F_ROH_ was not appreciably correlated with median reported height at the shoulder (Fig S1a, *P*=.71).

Body size has been observed to impact both age at first estrus, ovulation frequency, and parity across dog breeds (Borge et al, 2011). To ascertain whether body size was impacting litter size in this cohort, we regressed litter size against median shoulder height. We found a statistically insignificant positive association between median height at the shoulder and litter size (Fig S1b, *P*=.19).

Finally, age at time of parturition has been shown to impact litter size (Borge et al, 2011; Mandigers et al, 1994). We regressed litter size against the dog’s age at the time of litter recording and did not observe an appreciable correlation between these two factors (Fig S1c, *P*=.65).

The canine interestrus cycle is roughly seven months with high variation across breeds; bitches can also vary individually in their interestrus cycle depending on age and season (Sokolowski et al, 1977; Concannon et al, 1986; Davidson, 2006). Shorter interestrus periods, ergo, more frequent estrous cycles (heats), provide greater opportunities for conception and could therefore contribute to high conception rates. We plotted recorded annual heat frequency versus F_ROH_, separating samples by calendar age. We saw no significant correlation between estrous cycle frequency and F_ROH_ at any age. (Fig S2); however, we did note that dogs who had more than 1 heat per year were likely to maintain this higher than average heat frequency over all years recorded.

**Figure 2.**
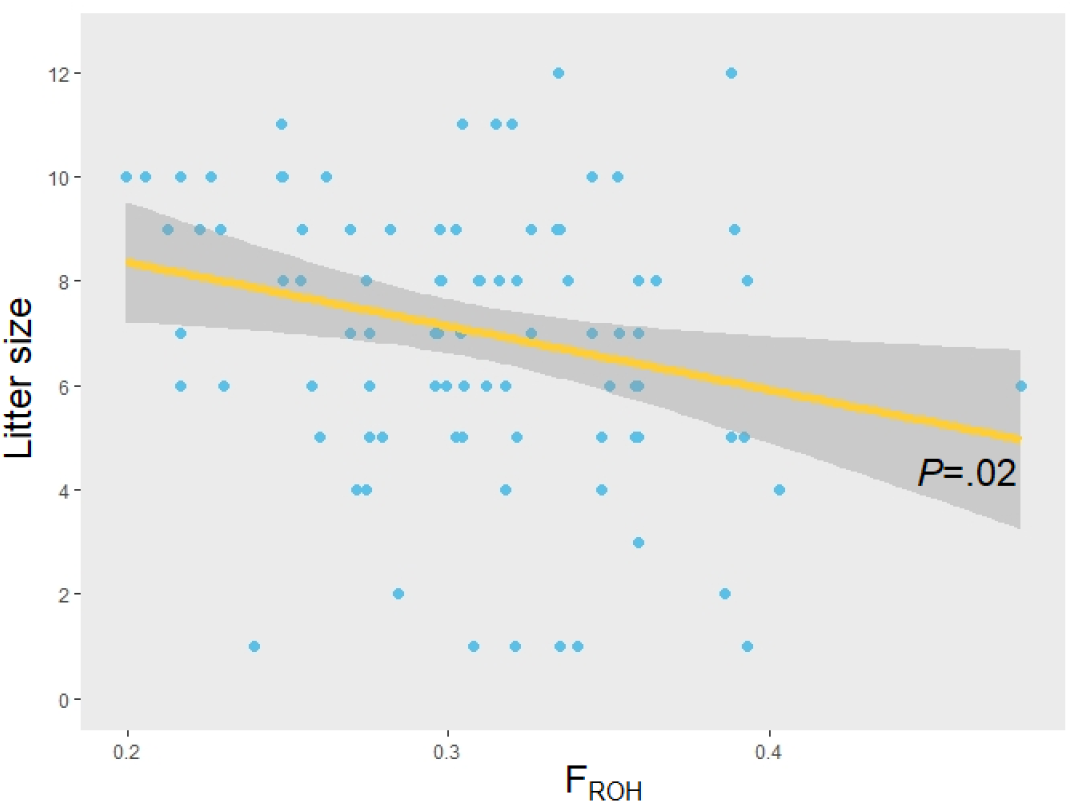
Litter size as measured by number of live puppies born is inversely correlated with F_ROH_. Individual litters are plotted in blue; linear regression (R^2^=0.102, *P*=.02) is shown in yellow with 95% confidence interval in gray.

We next measured the association of successful conception rate (SCR) versus F_ROH_. SCR is a derived value calculated from total number of litters produced over total number of attempted breedings. Dogs who had been bred one or less times were excluded from this analysis under the assumption that a single breeding (which would result in an SCR of either 0% or 100%) may not be reflective of a dog’s potential for SCR. We found that, while dogs with lower F_ROH_ had subjectively higher SCR, this result was not statistically significant (Fig S3).

We next regressed F_ROH_ against the number of live puppies born per litter using a mixed effects linear model, considering F_ROH_, median height, and age at time of litter log as fixed effect variables and dam ID as a random effect variable. We found a statistically significant negative correlation between F_ROH_ and number of live puppies (Fig 2, *P*=0.02).

A alternative mixed effects linear model was performed using F_ROH_, median height, and age at time of litter log as fixed effect variables and dam ID as a random variable, defining a standardized kinship matrix generated from GEMMA as the variance family to be used for the dam ID. This model also yielded a statistically significant negative correlation between F_ROH_ and number of live puppies (*P*=0.02).

While other measures of reproductive success could include variables for parturition and postnatal care, our dataset included just 5 reported cases of dystocia and 1 case of mastitis; data on puppy survival and progress post-partum were not available in all cases. However, post-natal measurements for reproductive success are likely to be much more complex in nature, and will likely require a much larger dataset to inform them.

## DISCUSSION

We and others have already demonstrated the potential of direct-to-consumer genomics to discover novel genetic variants affecting coloration (Deane-Coe et al, 2018; Eriksson et al, 2010), behavior (Hyde et al, 2016), and disease risk (Chang et al, 2017). Our present findings also emphasize the power of multi-institutional collaboration to expedite and improve the process of data-driven discovery. The longitudinal, all-encompassing nature of the GRLS represents a wealth of phenotypic data. Combined with high-quality, high-density SNP genotyping, the potential for rapid identification of genetic contributions to lifespan and healthspan in the Golden Retriever is unprecedented. The work described here is clear evidence: even with a relatively small sample size of purebred Golden Retrievers, we describe a statistically significant negative correlation between F_ROH_ and litter size.

The effects of inbreeding on reproductive success can be obscured by genotypic and phenotypic variation in the sample population. By using a subset of GRLS participants, we find ourselves in the lucky position of assessing this complex relationship in a natural population with, by definition, minimal variation. We do not observe a significant correlation between litter size and maternal body weight, though this has described by others (Borge et al, 2011). However, litter size trends have historically been documented across, but not within breeds, and it could be possible that body size variation within a breed with an already narrow range of acceptable body size could be insufficient to impact litter size. This hypothesis could be more definitively assessed in a larger sample set. Similarly, the negative effect of inbreeding on body weight has been explored in many species (reviewed in LeRoy et al, 2014). While we observe a subtle negative relationship between F_ROH_ and median shoulder height, in this cohort, this correlation was not significant, suggesting that a larger sample set could prove more informative.

Strikingly, the only variable that significantly impacts litter size in this cohort is F_ROH_. A negative correlation between pedigree-based estimates of inbreeding and litter size has been reported (LeRoy et al, 2015). To our knowledge, our work is the first to identify a significant correlation between a genomic estimate of inbreeding, F_ROH_, and fecundity, predicting a roughly one puppy reduction in litter size with every 10% increase in F_ROH_.

We also identify a suggestive negative correlation between successful conception rate, a measure derived from number of attempted breedings versus number of litters born. Given the many variables upon which successful conception depends upon, for example, appropriate timing of breeding relative to estrus, semen viability, and method of breeding, it is perhaps unsurprising that in this small cohort, this correlation was statistically insignificant. As such, we intend to examine SCR and other measures of fecundity in a larger cohort of Golden Retrievers. In addition, pending availability of phenotype, we would be eager to examine the effects of inbreeding on other indices of fertility including early fetal resorption, incidence of dystocia or perinatal complications, or, from the male point of view, sperm count or motility.

Purebred animal registries are no stranger to popular sire effect. Animals with significant titles and accomplishments are more likely to contribute to the next generation with the hopes that progeny will exhibit the same excellent performance, conformation, or work ethic of the parent. Perhaps the most dramatic example of popular sire effect exists within the Thoroughbred racehorse industry (Catton and Wezerek, 2018). However, selective use of just a few highly accomplished individuals essentially pushes the population into an artificial bottleneck, leading to reduced genetic diversity in the next generation. In the purebred dog world, certain measures do exist to control popular sire effect (Federation Cynologique Internationale, American Kennel Club (b)); further, most purebred dog breeders keep meticulous records in order to monitor and control the relatedness of their breeding animals. However, pedigree analysis of large populations of dogs still demonstrate a reduction in effective breeding population over the past 50 years (Calboli et al, 2008). Though our analyses remain preliminary, it is possible that the consequences of popular sire usage and the contribution of just a select number of individuals to the next generation have come to roost for many well-known dog breeds. We believe that this work sets the stage for a much larger population analyses by which regions of the genome associated with aspects of inbreeding depression--higher mortality, reduced reproductive success--could be pinpointed and breeding recommendations could be made to increase heterozygosity in these regions. In this regard, high density, high resolution genotyping could be invaluable for the maintenance and perpetuation of popular dog breeds.

## MATERIALS AND METHODS

Genomic DNA and phenotype information relative to reproductive status and success was requested from 100 female intact Golden Retriever dogs enrolled in the GRLS study had been bred at least once (Document S1).

Phenotype information was compiled and provided by the MAF; information was gathered via veterinary-and owner-submitted questionnaire annually and at each veterinary visit per MAF guidelines. Participants’ date of birth, physical exam findings, most recent estrous (heat) cycle and duration, date and method of last breeding and litter, litter size (puppies born, puppies weaned), and reproductive complications (dystocia, pyometra) were included.

Peripheral blood mononuclear cell (PBMC)-derived gDNA for each dog was provided by the MAF. gDNA was diluted to roughly 200 ng/uL; 50 uL of each sample as submitted for genotyping using on the Embark 220K SNP array platform as previously described (Deane-Coe et al, 2018). F_ROH_ was calculated using runs of homozygosity ≥ 500 kb as described in Sams et al, 2018. Successful conception rate (SCR) was calculated as the ratio of attempted breedings to number of litters born for each dog; dogs with zero attempted breedings were excluded from analysis. Violin plots of SCR relative to COI quartiles and regression plots for litter size relative to COI were generated with ggplot2 (Wickham, 2016).

Litter size was calculated as the variable livepup, number of live puppies born, compiled from MAF records. A linear mixed model (coi.with.barcode) was generated with the lmer function (lme4, Bates et al, 2015) in R, considering F_ROH_ (coi_with_public), median withers height (median_height), and age in years at the time of litter recording (age_at_visit_year) as fixed effect variables and unique dam ID (barcode) as a random effect variable (as described in Cnaan et al, 1998 and implemented in Lüpold et al, 2010, Koch et al, 2008):

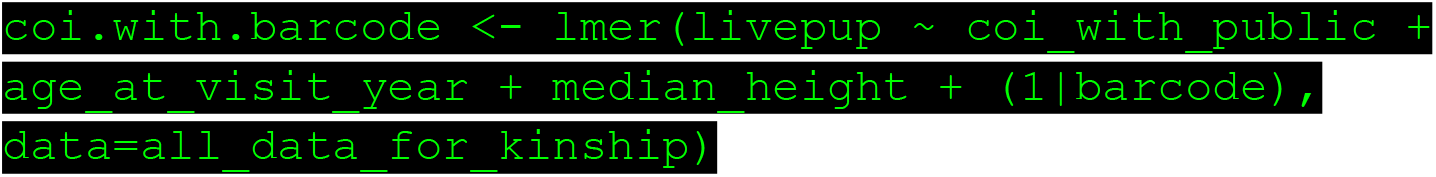

A second linear mixed-effects model (coi.with.kinship) was performed using lmekin function in coxme (Therneau 2018):

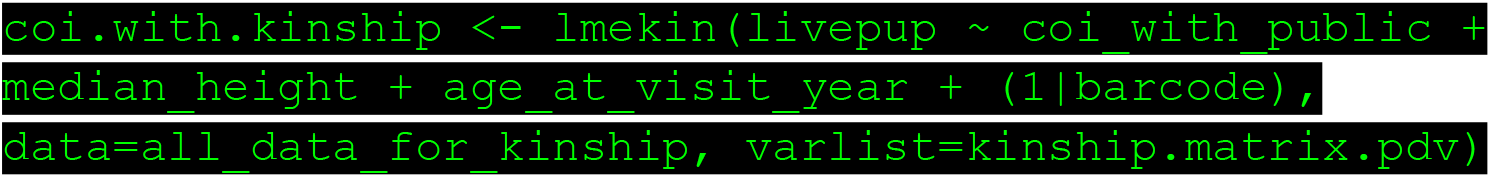

Using a standardized kinship matrix (kinship.matrix.pdv) generated with GEMMA (version 0.97) as a random effective variable (Zhou et al, 2012).

For all regressions, significance of Pearson’s correlation coefficient is reported as *P.*

## DATA AVAILABILITY

Additional data and complete summary statistics from the analyses in this paper will be made available to researchers through Embark Veterinary Inc., under an agreement with Embark that protects the privacy of Embark customers and their dogs. Please contact the corresponding author for more information and to apply for access to the data.

## ACKNOWLEDGEM ENTS

We thank the members of the Embark Discovery Team, Andrea Slavney, Taki Kawakami, Brett Ford, Sam Vohr, and Meghan Jensen for their suggestions and feedback on the contents of this manuscript. We thank Jackie Bubnell and Evan Buntrock for their insights into mixed linear modeling. We thank Brandon Goode for his discerning eye in manuscript finalization. And most importantly, we thank the Golden Retriever owners, breeders, and veterinarians of the Golden Retriever Lifetime Study: without your dedication, this work would not be possible.

## AUTHOR CONTRIBUTIONS

ETC analyzed the data and wrote the paper. AJS analyzed data. ARB and AJS jointly directed the research and writing. ARB wrote the proposal requesting samples to the MAF. MJ provided phenotype data and genomic DNA samples from MAF participants. KD and RP take management and leadership roles in the MAF and provide comments on the paper.

## AUTHOR INFORMATION

Conflict of interest statement: ETC, ARB, and AJS are employees of Embark Veterinary, a canine DNA testing company. ARB is co-founder and part owner of Embark. Correspondence and requests for materials should be addressed to PED, ARB, or AJS.

## SUPPLEMENTAL FIGURES & TABLES

**Fig S1.**
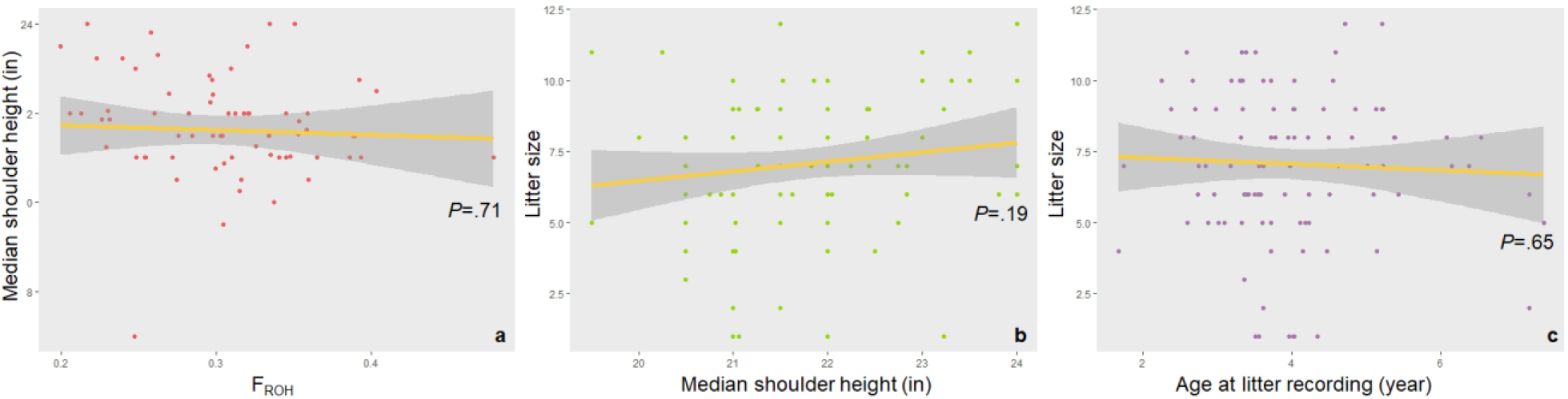
**Linear regression plots of a)** median shoulder height against F_ROH_, **b)** litter size against median height, and **c)** litter size against age at time of litter recording. While correlations between all three sets of factors have been demonstrated in the literature, we do not observe significant correlations between them in this dataset.

**Fig S2.**
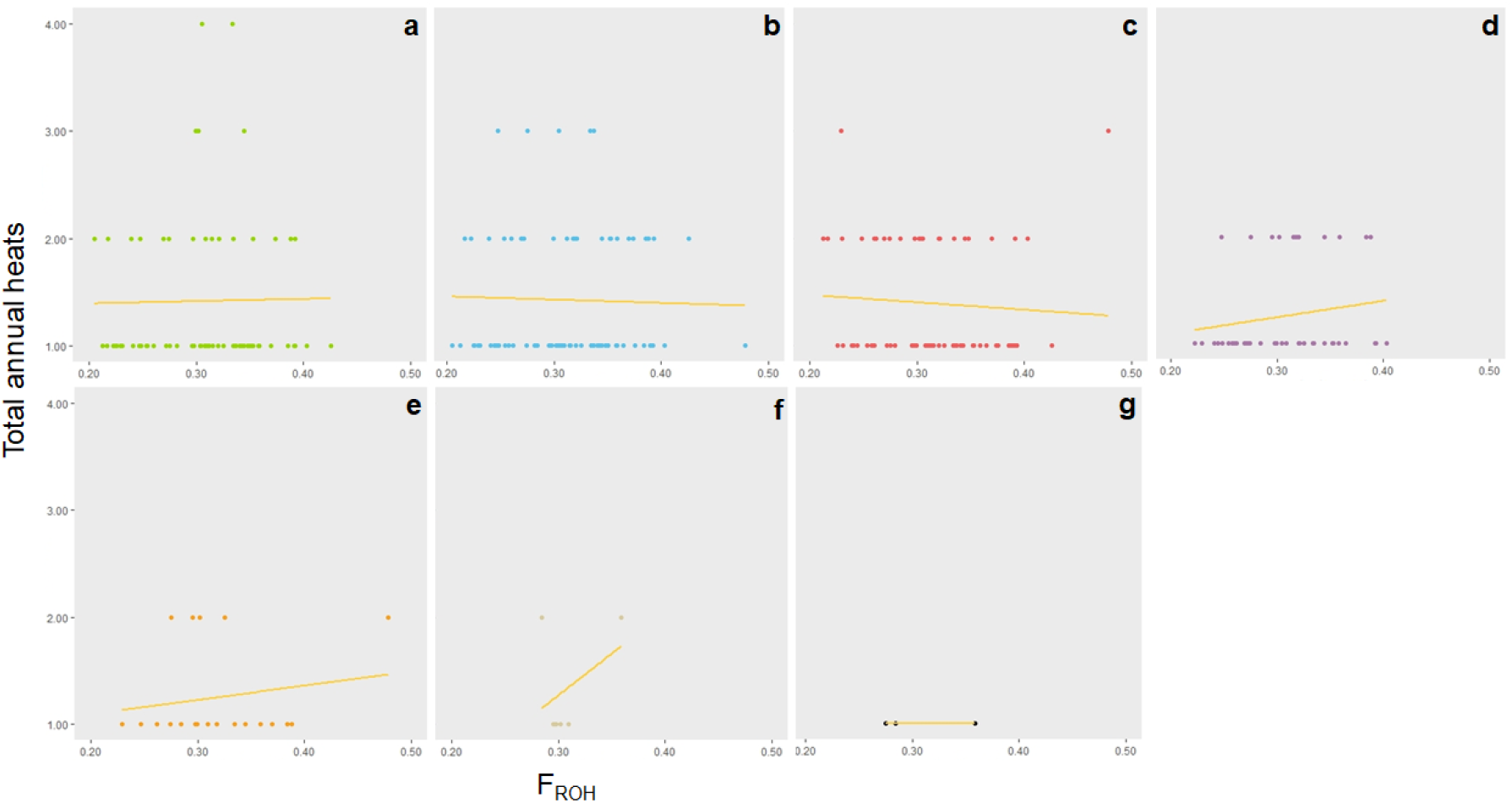
Regression of estrous cycles (heats) recorded against F_ROH_ for Embark-GRLS dogs at a) 1, b) 2, c) 3, d) 4, e) 5, f) 6, and g) 7 years of age. Recorded heats range from 1 to 4 heats per year at any age with no significant correlation between with F_ROH_ and annual heats at any year of age. Data points decrease dramatically for 6 (tan) and 7 (black) year old dogs.

**Fig S3.**
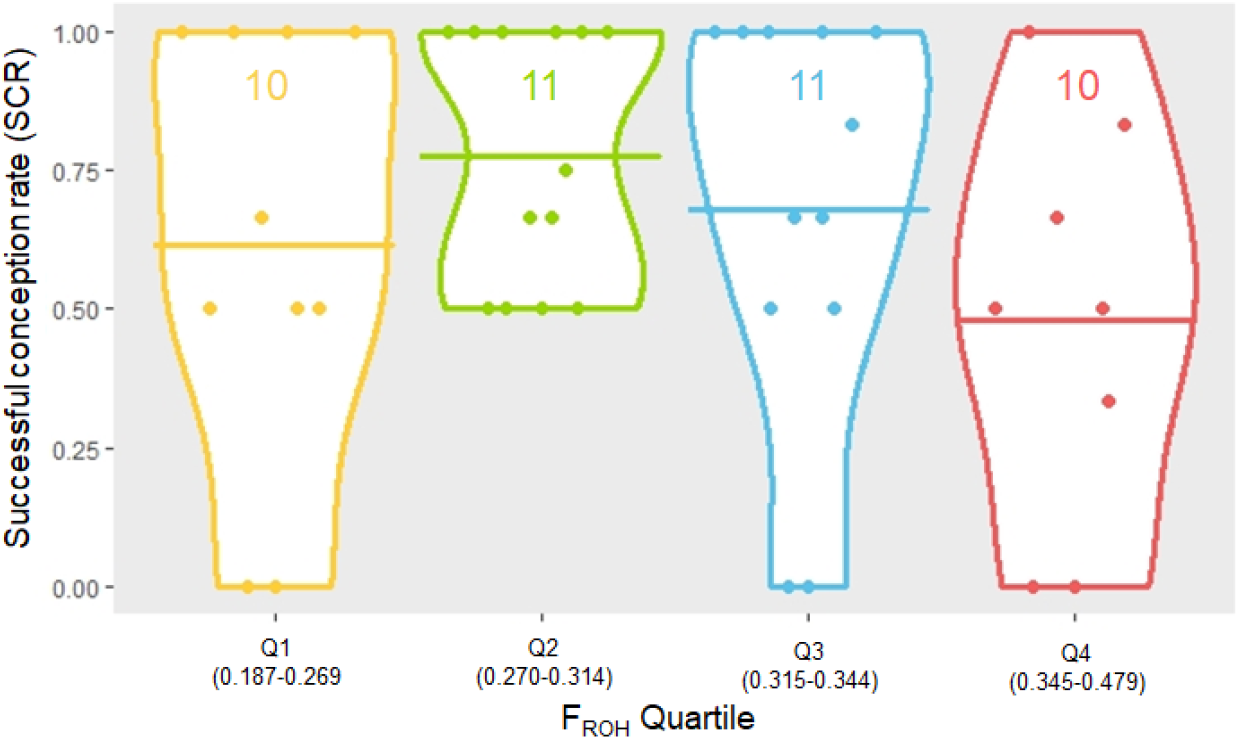
Violin plots of SCR of GRLS dogs who had been bred at least twice (n=44) separated into F_ROH_ quartiles. Number of observations per quartile are indicated in each violin plot. Crossbars represent average SCR within quartiles Dogs in the fourth quartile (0.344 < F_ROH_ < 0.479) have a lower average SCR than dogs in the first, second, and third quartiles, though this difference is insignificant (*P*=.21).

**Table S1:** Summarizing table of statistics on the 100 dogs selected for the Embark-GRLS cohort. Note that not all dogs represented in this table were genotyped; their statistics are not included in this study.

| Characteristic | Integer Age (years) |  |  |  |  |  |  |
| --- | --- | --- | --- | --- | --- | --- | --- |
|  | 1 | 2 | 3 | 4 | 5 | 6 | 7 |
| Number of visits | 63 | 133 | 136 | 109 | 46 | 19 | 6 |
| Median body condition score (min, max) | 4(4,7) | 4(2,7) | 5(4,7) | 5(2,6) | 5(2,8) | 5(3,7) | 5(5,7) |
| Mean Height at shoulders in cm (±sd) | 54.7(3.2) | 55.2(3.2) | 54.9(3.1) | 54.6(2.6) | 54.9(2.3) | 53.8(1.5) | 54.6(2.0) |
| Mean Weight in kg (±sd) | 25.4(3.8) | 26.7(3.5) | 27.9(3.4) | 28.3(3.2) | 27.6(3.7) | 28.8(4.5) | 28.9(4.8) |
| Total breedings | 1 | 13 | 58 | 62 | 27 | 8 | 3 |
| Total litters | 0 | 7 | 41 | 42 | 20 | 5 | 3 |
| % of breeding resulting in litter | 0% | 54% | 71% | 68% | 74% | 63% | 100% |
| Total live births (min, max) | n/a | 59(4,11) | 273(2,11) | 251(2,10) | 143(2,12) | 32(7,10) | 13(2,6) |
| Total stillborn (min, max) | n/a | 3(0,3) | 24(0,3) | 33(0,5) | 7(0,2) | 1(0,1) | 2(0,2) |
| Total weaned (min, max) | n/a | 56(4,10) | 227(0,10) | 241(1,10) | 125(2,12) | 32(7,10) | 13(2,6) |
| Number of c-sections<br>(% of pregnancies) | n/a | 2(28%) | 13 (32%) | 9(21%) | 4(20%) | 0 | 0 |

